# Screening anti-infectious molecules against *Mycobacterium ulcerans*: a step towards decontaminating environmental specimens

**DOI:** 10.1101/2020.03.31.018382

**Authors:** N Hammoudi, R Verdot, J Delorme, A Bouam, M Drancourt

**Author notes:** Corresponding author: Prof. Michel DRANCOURT, IHU Méditerranée Infection, MEPHI, 19-21 Boulevard, Jean Moulin, 13005 Marseille, France. mailto.

## Abstract

*Mycobacterium ulcerans*, a non-tuberculous mycobacterium responsible for Buruli ulcer, is residing in poorly defined environmental niches in the vicinity of stagnant water points where very few isolates have been confirmed. In the perspective of culturing *M. ulcerans* from such contaminated environmental specimens, we tested the *in vitro* susceptibility of *M. ulcerans* CU001 strain co-cultivated with XTC cells to anti-infectious molecules registered in the French pharmacopoeia, using a standardised concentration, to find-out molecules inactive against *M. ulcerans* which could be incorporated in decontaminating solution. Of 116 tested molecules, 64 (55.1%] molecules including 34 (29.3%] antibiotics, 14 (12%] antivirals, 8 (6.8%] antiparasitic and 8 (6.8%] antifungals were ineffective against *M. ulcerans* CU001; leaving 52 molecules active against *M. ulcerans* CU001. Three such inactive antimicrobial molecules (oxytetracycline, polymyxin E and voriconazole] were then selected to make a decontamination solution shown to respect *M. ulcerans* CU001 viability. These three antimicrobials could be incorporated into a decontamination solution for the tentative isolation and culture of *M. ulcerans* from environmental samples.

## Introduction

*Mycobacterium ulcerans* is a nontuberculous mycobacterium responsible for Buruli ulcer, an opportunistic neglected tropical disease that also affects some nonhuman mammal species [1]. *M. ulcerans* was first isolated in Australia in 1948 after the disease was initially described in 1897 in Uganda [2]. Phylogenetic analysis showed that *M. ulcerans* evolved from a common ancestor with *Mycobacterium marinum* after genomic reduction characterized by an accumulation of insertion sequences and counterbalanced by the acquisition of a giant plasmid encoding for the nonribosomal synthesis of mycolactones, exotoxins exhibiting ulcerative, analgesic, immunosuppressive and anti-inflammatory properties [3–4]. *M. ulcerans* is an environmental mycobacteria, and although DNA sequences specific to *M. ulcerans* are routinely detected in aquatic ecosystems by PCR [5–6], its exact reservoir and routes of transmission to humans remain unknown [7]. Indeed, PCR-based data do not provide insight into the viability of these detected mycobacteria.

The first environmental *M. ulcerans* isolate was reported in 2008 from an aquatic insect [8], 60 years after it was first isolated from a patient [2]. The long delay between isolation from environmental sources and clinical sources illustrates the particular constraint isolating *M. ulcerans* from environmental sources, i.e., contamination by fast-growing mycobacteria, bacteria and fungi [9–10–11]. Therefore, an efficient decontamination protocol is key for the tentative isolation of environmental *M. ulcerans* strains. From this perspective, different strategies have been used, including the Petroff and reversed Petroff methods, the combinations of oxalic acid-NaOH, NaOH-malachite green-cycloheximide, and N-acetyl-cysteine-NaOH, and Löwenstein-Jensen (LJ] medium supplemented with polymyxin B, amphotericin B, nalidixic acid, trimethoprim and azlocillin (PANTA], mycobactin J, isoniazid and ethambutol [1]. However, all these decontamination methods can adversely affect the viability of *M. ulcerans* [12].

To progress towards an efficient decontamination method that preserves the viability of *M. ulcerans*, we tested the nonantimicrobial activity of 116 antimicrobial agents, including antibiotics, antifungals, antiparasitic and antiviral drugs listed in the French pharmacopeia, against *M. ulcerans* to identify molecules that could potentially be used to isolate environmental *M. ulcerans* strains using xenic and axenic culture media.

## Materials and methods

### Antimicrobial drugs

A collection of 116 molecules registered as antimicrobials in the French pharmacopoeia were tested for their activity against *M. ulcerans*. Precisely, 70 molecules were purchased from pharmaceutical industries, 45 molecules from Sigma (Lezennes, France] and one from EUROMEDEX (Souffelweyersheim, France] (supplementary Table. 2]. Each molecule was resuspended in the appropriate solvent according to the supplier to prepare a stock solution that was aliquoted and stored at −20°C.

### Mycobacteria and cells

*M. ulcerans* CU001, a clinical isolate from a Ghanaian patient, was kindly provided by Professor Vincent Jarlier, Salpêtrière hospital, Paris France [13], and cultured in a biosafety level 3 laboratory on Middlebrook 7H10 agar (Becton Dickinson, Le Pont de Claix, France] supplemented with 10% oleic acid, bovine albumin, dextrose and catalase enrichment (OADC, Becton Dickinson] for six weeks under an aerobic atmosphere at 30°C. Colonies were suspended on a 20-mL glass tube containing glass beads and 10 mL of Dulbecco’s phosphate buffered saline (DPBS]. The suspension was vigorously vortexed and passed three times through a 26-gauge needle to separate the aggregates. The suspension was then calibrated to 10^6^ colony forming units (CFUs]/mL corresponding to 1 Mcfarland. In parallel, XTC cells originating from the South African clawed toad *Xenopus laevis* [14] were grown in L15 glutaMAX medium (Gibco, ThermoFisher, Illkirch, France] supplemented with 5% heat-inactivated fetal bovine serum (FBS] and 40 mL of tryptase in 75 cm^2^ flasks at 28°C for 7 days. Cells were detached by tapping the flasks, and 2 mL of cell suspension was transferred to each well of a 12-well cell culture plate (ThermoFisher]. In this study, XTC cells were used as a support for cultivating *M. ulcerans* and to test the activity of molecules to mimic the biotic environment in which *M. ulcerans* is suspected to thrive.

### Antimicrobial assay

The nonactivity of each of the 116 antimicrobials investigated against *M. ulceran*s was assayed in a coculture system with XTC cells and *M. ulcerans*. In brief, each 12-well plate was divided into four columns of three wells. Then, 20 μL of DPBS was added to each well in column one (negative control], 20 μL of *M. ulcerans* CU001 suspension at 10^6^ CFUs/mL (equivalent to 0.2 × 10^5^ CFUs per well] was added to each well in column two (positive control], and 20 μL of the same *M. ulcerans* inoculum was added to each well in the third and fourth columns, which were supplemented with antibiotics, one antibiotic per column. The final concentration of each antibiotic used in these experiments was five times the minimum inhibitory concentration (MIC] reported in the literature. The plate was incubated at 30°C under an aerobic atmosphere for seven days; then, 100 μL of each well was plated on Middlebrook 7H10 agar plates in triplicate and incubated at 30°C for six weeks. The number of colonies (up to 150 colonies] on each plate was counted using Fiji-ImageJ software (https://imagej.net/Fiji/Downloads], and the average number of colonies grown on the three plates was calculated for each antibiotic. In this study, any molecules that left > 75 colonies on plates (50%] were considered to not affect the survival of *M. ulcerans*.

### Trans *Mycobacterium ulcerans* (Trans MU], a decontamination medium for the recovery of *M. ulcerans*

We selected three antimicrobial agents that target three categories of contaminating organisms that were previously encountered when attempting to grow *M. ulcerans* from environmental samples (Bouam A., Hammoudi N, unpublished data], i.e., an anti-*Bacillus* compound (oxytetracycline], an anti-gram-negative bacteria compound (polymyxin E] and an antifungal agent (voriconazole] at the same concentrations that were used in this study (Table.1]. In the first step, the antimicrobial mixture was mixed with 1% chlorhexidine (final concentration] to create a decontamination medium (Trans MUl]. In the second step, the antimicrobial mixture was mixed with Middlebrook-OADC at 50°C and poured into 50-55-mm Petri dishes to create a decontamination and culture agar medium (Trans MUg]. A 10^5^ CFUs/mL suspension of *M. ulcerans* was prepared in sterile phosphate buffered saline (PBS], which was previously used as a negative control for *M. ulcerans* DNA in an RT-PCR experiment. A 4-mL volume of TRANS MUl was added to 1 mL of *M. ulcerans* suspension and incubated for 4 days at room temperature with constant shaking. Then, the mixture was centrifuged for 15 min at 3,500 g, and the pellet was washed and vortexed vigorously in 5 mL of a neutralizing solution (composed of 200 mL of PBS, 0.6 g of egg-lecithin and 2 mL of Tween 80] [15]. The suspension was centrifuged at 3,500 g for 15 min, and the pellet was resuspended in 1 mL of sterile PBS. A 200-μL volume of sample was inoculated in triplicate on Trans MUg at 30°C for 2 months. In parallel, a sample of sterile PBS contaminated with 10^5^ CFUs/mL *M. ulcerans* was directly plated on Middlebrook 7H10 agar and TRANS MUg in triplicate. A second sample treated only with TRANS MUl was plated on Middlebrook 7H10 agar in triplicate as a control.

**Table 1.** Results of the sensitivity of *M. ulcerans* against 116 antimicrobials, the color gradient from red to yellow includes antimicrobials that affect the viability of *M. ulcerans* CU001. The color gradient from yellow to green includes antimicrobials that do not affect the viability of *M. ulcerans*. (cut of 50% = 75 Avg CFU Nbr].

| Anitmicrobials | N° CFUs average | Anitmicrobials | N° CFUs average | Anitmicrobials | N° CFUs average | Anitmicrobials | N° CFUs average |
| --- | --- | --- | --- | --- | --- | --- | --- |
| Aztréonam | 0 | Kanamycine | 10,66 | Famciclovir | 90 | Ornidazole | 150 |
| Cefalexine | 0 | Tigécycline | 16,33 | Sulfadiazine | 94,33 | Oxytétracycline | 150 |
| Céfépime | 0 | Sulfamethoxazole | 17 | Colistine | 103 | Péfloxacin | 150 |
| Céfotaxime | 0 | Acide fusidique | 17,66 | Luméfamtrine | 109,5 | Sulbactam | 150 |
| Céfradine | 0 | Fumagilline | 17,66 | Zanamivir | 117,33 | Témocilline | 150 |
| Daptomycine | 0 | Chloramphenicol | 18,33 | Nétilmicine | 120 | Thiamphenicol | 150 |
| Ertapénem | 0 | Pipéracilline/tazobactam | 26,33 | Piperaquine | 127 | Isoniazide | 150 |
| Érythromycine | 0 | Halofantrine | 28,33 | Voriconazole | 129,66 | Griséofulvine | 150 |
| Gentamicine | 0 | Spiramycine | 28,66 | Fluconazole | 136,66 | Itraconazole | 150 |
| Gramicidine | 0 | Oxacilline | 31,66 | flubendazole | 140 | Micafungine | 150 |
| Lévofloxacin | 0 | Doripénem | 32,33 | Atovaquone | 146 | Terbinafine | 150 |
| Lincomycine | 0 | Penicilline G | 36,33 | Fosfomycine | 147 | Caspofungine | 150 |
| Nitrofurantoïne | 0 | Tinidazole | 40 | Ampicilline | 150 | Niclosamide | 150 |
| Rifabutine | 0 | Secnidazole | 42,66 | Azithromycine | 150 | Pentamidine | 150 |
| Tétracycline | 0 | Bacitracine | 47,66 | benzylpénicilline | 150 | Praziquantel | 150 |
| Triméthoprime | 0 | Teicoplanine | 50 | Céfaclor | 150 | Pyrantel | 150 |
| Vancomycine | 0 | Artemether | 51,33 | Céfixime | 150 | Pyrimethamine | 150 |
| Chloroquine | 0 | Mitomycine C | 59,66 | Ceftaroline | 150 | Aciclovir | 150 |
| Méfloquine | 0 | Pipérazine | 60 | Ceftazidime | 150 | Amantadine | 150 |
| Rifampicine | 0,66 | Tobramycine | 61,66 | Céfuroxime | 150 | Cidofovir | 150 |
| Cefadroxil | 1 | Augmentin | 65 | Cepfodoxim | 150 | Foscarnet | 150 |
| Métacycline | 1 | Entécavir | 71,66 | Clindamycine | 150 | Oseltamivir | 150 |
| Proguanil | 1 | Spectinomycine | 74,33 | Cloxacilline | 150 | Ritonavir | 150 |
| Triclabendazole | 1 | Valaciclovir | 76 | Imipénème/cilastatine | 150 | Telbivudine | 150 |
| Pyrazinamide | 1,66 | Ceftriaxone | 77 | Josamycine | 150 | Ténofovir | 150 |
| Doxycycline | 2 | Polymyxine B | 78 | Méropénem | 150 | Valganciclovir | 150 |
| Clofazimine | 3 | Cefazoline | 79 | Métronidazole | 150 | Flucytosine | 150 |
| Céfotiam | 5 | Amphotéricine B | 84 | Mupirocine | 150 | Ganciclovir | 150 |
| Sulfadoxine | 5,66 | Ethambutol | 84,66 | Néomycine | 150 | Lamivudine | 150 |

## Results

### Antimicrobial assay

A total of 116 pharmaceutical molecules corresponding to four therapeutic families were tested against *M. ulcerans* CU001. As a result, 64 (55.2%] molecules did not affect the viability of *M. ulcerans*; these molecules comprised 34 (29.3%] antibiotics, 14 (12%] antivirals, 8 (6.8%] antiparasitics and 8 (6.8%] antifungals, as displayed in cells colored yellow to green in (Supplementary Table 1.]. In contrast, 52 (44.8%] molecules altered the viability of *M. ulcerans*; these molecules comprised 42 (36.2%] antibiotics and 9 (7.6%] antiparasitics in addition to entecavir, which yielded 71 colonies, just below the 75-colony cut-off used in this study, while no other antivirals or antifungals altered the viability of *M. ulcerans*, as displayed in cells colored in red to yellow in (Supplementary Table 1.].

### Trans MU a decontamination medium

A *M. ulcerans* suspension in PBS was used as a positive control and yielded colonies after 30 days of incubation at 30°C. The same observation was reported for PBS contaminated with 10^5^ CFU/mL *M. ulcerans* cultivated on TRANS MUg, which was positive after 30 days of incubation at 30°C. In addition, Middlebrook 7H10 agar plates or TRANS MUg plates inoculated with 10^5^ CFU/mL *M. ulcerans* previously treated with TRANS MUl allowed *M. ulcerans* to grow after 30 days of incubation at 30°C.

## Discussion

We are presenting the first, large, open-minded study of the activity of 116 molecules against *M. ulcerans*, the etiologic agent of Buruli ulcer [1]. The data reported herein were validated by the positivity of positive controls, and the results reported here are in agreement with those previously published in the literature, such as the *in vitro* activity of rifampicin against *M. ulcerans*, which is the current basis of Buruli ulcer treatment [16–17]. and with the In *silico* prediction of genes that confer resistance to isoniazid and pyrazinamide [18], The data reported here were obtained by incorporating XTC cells in anti-*M. ulcerans* assays; the XTC cells were used as surrogates of the organic environments naturally encountered by *M. ulcerans* in its ecological niches, as well as in clinical tissues in which *M. ulcerans* behave as a pathogen.

This study broadens the spectrum of molecules active against *M. ulcerans in vitro*, including 20 previously reported antibiotic molecules [1] and 50 molecules encompassing not only antibiotics but also antiparasitics and an antiviral (entecavir]; the MICs of these molecules are still pending determination, a task beyond the spectrum of the present study. Interestingly, this study shows that subtle chemical differences, such as tetracycline oxidation to oxytetracycline, may modify the activity of molecules against *M. ulcerans*. Pursuing this type of observation was beyond the spectrum of this study, but data reported herein could be used for the chemical design of molecules active against *M. ulcerans*. This will be of interest because Buruli ulcer is classified as a neglected tropical infection [19].

The original aim of this study was to identify molecules inactive against *M. ulcerans* in such a way that they could be incorporated into decontamination solutions, while maintaining the viability of the pathogen; these decontamination solutions could be used in the medical field for the transportation of naturally contaminated ulcer clinical samples collected to identify *M. ulcerans* or for the examination of decontaminated environmental samples collected in ecological niches where *M. ulcerans* had been detected by PCR-based methods [20]. Indeed, decontaminating or removing contaminating microorganisms from clinical and environmental samples is an essential step to isolating *M. ulcerans* [9]. From this perspective, we identified 66 inactive molecules against *M. ulcerans in vitro*; these different molecules may therefore be potential antimicrobials to be incorporated into culture media to decontaminate environmental and clinical samples for the isolation of *M. ulcerans*. Several studies have reported that delays in sample transport may affect the viability of *M. ulcerans* [21]. In addition, several decontamination methods proposed for the isolation and culture of *M. ulcerans,* such as the Petroff method, the reversed Petroff method and the oxalic acid decontamination methods [22], or the use of HCl, all reduce *M. ulcerans* viability resulting in culture failure [9]. However, commercially available mixture PANTA (Becton Dickinson] does not modify *M. ulcerans* growth contrary to what has been reported for *Mycobacterium leprae* [23].

In the present study, we developed a protocol that allowed *M. ulcerans* subculture after only four days of preculture in TRANS MUl followed by 30 days of incubation on TRANS MUg at 30°C. This protocol is now used for the tentative isolation and culture of *M. ulcerans* from environmental samples, including aulacode feces samples in which *M. ulcerans* has been previously isolated but not subcultured [24].

## Supplementary material legends

**Supplementary Table 1**. Classification of the 116 antimicrobials used in this study according to their pharmaceutical class and the sensitivity of *M. ulcerans* CU001.

**Supplementary Table 2**. Source of 116 molecules registered as antimicrobials in the French pharmacopoeia and used in these studies.

## Acknowledgments

This work was supported by Région Provence Alpes Côte d’Azur

